# Identification and characterization of alamandine-(1-5), a new component of the Renin-Angiotensin System with unique properties

**DOI:** 10.1101/2024.04.27.591083

**Authors:** Melissa Tainan Silva Dias, Sthefanie Chaves de Almeida Gonçalves, Filipe Alex da Silva, Lucas Rodrigues-Ribeiro, Kamylle Silva Ferraz, Sérgio Scalzo, Matheus F. Itaborahy, Nícia Pedreira Soares, Danilo Augusto Alves Pereira, Pedro Alves Soares, João Batista Rodrigues Dutra, Carolina Fonseca de Barros, Uri Flegler Vieira-Machado, Isadora Zhong Liang Ferreira Feng, Ana Caroline Ventris de Godoy, Adelson Héric Alves Monteiro, Marcos Eliezeck, Bruno Sanches, André Monteiro, Amanda de Sá Martins de Bessa, Ana Paula Davel, Natália Nóbrega, Júlia Rezende Ribeiro, Maria Luiza Dias-Pinto, Bruno Durante da Silva, Leandro Eziquiel de Souza, Amanda de A. Silva, Michael Bader, Natália Alenina, Luciano dos Santos Aggum Capettini, Maria José Campagnole-Santos, Thiago Verano-Braga, Marco Antônio Peliky Fontes, Andrea Siqueira Haibara, Daniel Campos Vilella, Maria Claudia Irigoyen, Fernanda Ribeiro Marins, Carlos Henrique de Castro, Ana Cristina Simões-e-Silva, Maria de Fátima Leite, Silvia Guatimosim, Robson A. S. Santos

## Abstract

The renin-angiotensin system (RAS) comprises a biochemical cascade that hydrolyzes angiotensinogen into several different bioactive peptides, which can activate different receptors promoting plenty of specific effects. The aim of this study was to evaluate the presence of the putative product of alamandine, the pentapeptide alamandine-(1-5) in the circulation and its biological activity. To accomplish this we have used mass spectrometry (MALDI/TOF/TOF, LC-MS/MS) and several methodologies including isolated blood vessels, isolated perfused hearts, isolated cardiomyocytes, blood pressure recording in freely-moving normotensive and hypertensive rats (SHR), high resolution echocardiography (VEVO 2100), central administration (ICV infusion and microinjection in the insular cortex), cell culture (endothelial cells and GPCR-transfected CHO cells) and wild type and Mas, MrgD or AT2R deficient mice. Our results show that alamandine-(1-5) circulates in the human and rodent blood and promotes many biological central and peripheral actions. More importantly, its plasma concentration is increased in pediatric nephropathic patients. A major role for plasma ACE activity in the formation of alamandine-(1-5) from alamandine was observed using plasma samples from Angiotensinogen-KO mice. Alamandine-(1-5) increased Baroreflex sensitivity and produced a long-lasting (∼6 hours) anti-hypertensive effect in SHR, associated with a significant reduction in cardiac output. A particularly important effect of this pentapeptide was observed in isolated perfused heart and cardiomyocyte contractility (reduced inotropism). It was capable of stimulating NO production through all receptors from the renin-angiotensin protective arm, (MAS, MrgD and AT2R) in CHO-transfected cells. Our data shows that Alamandine-(1-5) exhibits selective actions that set it apart from traditional concepts of the vasodilatory axis of the RAS and that are possibly intricately linked to a complex interplay between Mas, MrgD and AT2 receptors. This novel finding suggests that RAS may possess a complexity that surpasses our current understanding.

## INTRODUCTION

It is well known that the renin-angiotensin system (RAS) plays a critical role in cardiovascular control and hydro-electrolyte homeostasis. The biologically active end products of this system are formed by a limited proteolysis process starting with the hydrolysis of the glycoprotein angiotensinogen by renin. Angiotensin(Ang) II and Ang-(1-7) are key players of the RAS acting as mediators of the classical and the non-canonic axes, respectively ^1^. Both peptides can undergo decarboxylation of the N-terminal amino acid aspartate to alanine, forming Ang A and alamandine, respectively. While Ang A displays actions that resemble those of Ang II, acting on AT1 and AT2 receptors, alamandine differs from Ang-(1-7) by acting on MrgD instead of Mas receptors ^2^. In addition, at least in some circumstances alamandine can have selective actions such as reported in the insular cortex ^3^. Hydrolysis of Ang-(1-7) by ACE can form the pentapeptide Ang-(1-5) which has been reported as an AT2R or Mas agonist depending on the cell type and organ^4–6^.

There is no data in the literature addressing the formation and actions of alamandine-(1-5) the putative product of alamandine through ACE hydrolysis. Here, we aimed to evaluate the formation and actions of the heptapeptide, alamandine-(1-5). Our results suggest that alamandine-(1-5) can be formed from alamandine by ACE and is present in the human blood. This novel angiotensin possesses unique activities including a marked effect in cardiac inotropism. These findings will contribute to importantly extend our understanding of the renin-angiotensin system.

## MATERIALS AND METHODS

### Drugs

Alamandine was purchased from RS synthesis (Louisville, KY USA). Alamandine-(1-5) was purchased from Byosynthan (Berlin-Buch, Germany). Angiotensin II and Angiotensin-(1-7) were from Bachem Basel, Switzerland). L-NAME was from Sigma-Aldrich (St. Louis, Missouri, United States). All other solvents and chemicals were reagent grade or equivalent. The suppliers of the other drugs and reagents were listed in each method.

### Experimental protocols procedures

All experiments involving animal were performed according to protocols approved by the Institutional Animal Care and Use Committee at Universidade Federal de Minas Gerais (UFMG). The study followed the National Institutes of Health (NIH) Guide for the Care and Use of Laboratory Animals.

### Mouse aortic rings

#### Animals

Male C57BL/6J, MrgD KO, AT2R KO, and MASR KO mice (10-14 weeks old) were obtained from the animal facilities of the Federal University of Minas Gerais. The animals were housed in a controlled temperature environment (25±2°C) with a 12 h light/dark cycle and a Standard diet and water were supplied *ad libitum*. All experimental protocols were approved and performed following the Animal Use Ethics Commission (CEUA-UFMG; protocol numbers 189/2023; 253/2022; 5/2018).

#### Isolated aortic ring preparation

Isolated aortic rings from the descending thoracic aorta were placed in the organ bath with Krebs-Henseleit solution (110.8 mmol.L^-1^ NaCl, 5.9 mmol.L^-1^ KCl, 25 mmol.L^-1^ NaHCO_3_, 1.07 mmol.L^-1^ MgSO_4_, 2.49 mmol.L^-1^ CaCl_2_, 2.33 mmol.L-1 NaH_2_PO_4_, 11.51 mmol.L^-1^ glucose; pH 7.4) contained 95 % O_2_ and 5 % CO_2_ at 37 °C, under a tension of 0.5 g for 60 minutes to stabilization. After the stabilization, the rings were pre-constricted with phenylephrine (10^-7^ mol.L^-1^) and received a cumulative concentration (10^-11^ to 10^-6^ nmol.L^-1^ mol.L^-1^) of alamandine-(1-5) to evaluate the vascular effect of this peptide. Some vessel segments were incubated with L-NAME (N-nitro-L-arginine methyl ester 100 μmol.L^-1^, Sigma-Aldrich), The mechanical activity was isometrically recorded using a force transducer and a data acquisition system (DATAQ Instruments, USA).

#### Isolated microvessels

Second order mesenteric resistance arteries (internal diameter <300 µm) were dissected out and mounted in a wire multichannel myograph (Model 620M, DMT A/S, Aarhus NA, Denmark) containing warmed (37 °C) and aerated (95% O_2_ and 5% CO_2_) physiological salt solution (in mmol.L^-1^: 118 NaCl, 4.7 KCl, 25 NaHCO_3_, 2.5 CaCl_2_.2H_2_O, 1.2 KH_2_PO_4_, 1.2 MgSO_4_.7H_2_O, 11 glucose, and 0.01 EDTA) ^7^. After a 30-minutes equilibration period, arteries were exposed to 60 mM KCl to determine the maximal tension. After washout and 30 min of equilibration, 1 μmol.L^-1^ phenylephrine (Sigma-Aldrich) was administered to induce contraction corresponding to the half-maximum KCl contraction, and then concentration-response curves to alamandine-(1-5) (10^-12^ to 10^-6^ mol.L^-1^) were determined.

### Isolated Adult Cardiomyocytes

#### Animals

Eight to 10-week-old male C57BL/6 mice were used in this preparation. All experiments were performed according to protocols approved by the Institutional Animal Care and Use Committee at Universidade Federal de Minas Gerais (UFMG). The study followed the National Institutes of Health (NIH) Guide for the Care and Use of Laboratory Animals.

#### Adult ventricular myocyte isolation

Briefly, hearts from 10-week old male mice were rapidly collected, conditioned into cold Tyrode solution (in mmol.L^-1^) 140 NaCl, 5 KCl, 1 MgCl_2_, 0.33 NaH_2_PO_4_, 1.8 CaCl_2_, 10 glucose, and 10 HEPES; pH 7.4 adjusted with NaOH), while the ascending aorta was cannulated and properly sealed. Ventricular myocytes were then isolated through the Langendorff method, which consists in an initial enzymatic dissociation, followed by mechanical dissociation, both at 37 °C. Freshly isolated ventricular myocytes were stored in typical Tyrode solution as previously described ^8^. Subsequently, cardiomyocytes were incubated for 10 min with alamandine-(1-5) (10^-7^ -10^-9^ mol.L^-1^), and intracellular Ca^2+^ [Ca^2+^]_i_ and cellular contractility were evaluated.

#### Ca^2+^ imaging and contractility recordings in ventricular myocytes

Intracellular Ca^2+^ [Ca^2+^]_i_ measurements were performed in cardiomyocytes loaded with Fluo-4 AM dye (5 μmol.L^-1^, ThermoFisher, Cat #F14201). After a 30-minute loading period, the cells were washed once with Tyrode at room temperature to remove excess dye. In sequence, the cells were treated with alamandine-(1-5), and stimulated to contract at 1Hz at room temperature. Imaging for intracellular Ca^2+^ concentration ([Ca^2+^]_i_) was acquired by using the Zeiss 880 confocal microscope (located at CAPI/UFMG), following established procedures as described before ^9,10^. For contractility measurements ventricular myocytes were kept in Tyrode’s solution, subjected to electrical stimulation using platinum electrodes, with a pulse frequency of 1 Hz and a voltage of 40 V, each pulse lasting 5 ms. To capture cellular images (resolution of 640x480 pixels, featuring a pixel size of 0.25 μm/pixel and 8-bit depth), a high-speed digital CMOS camera (SILICON VIDEO 642 M, EPIX, Inc) was utilized, operating at a frame rate of 200 fps. The data from cellular contractility was obtained according to the protocol previously described ^11^. After recording cycles of contraction-relaxation from untreated (control) cells, the myocytes were exposed to alamandine-(1-5) and contractility measurements were recorded again. Image processing and analysis were conducted using the CONTRACTIONWAVE software, employing the dense optical flow method ^12^.

#### Isolated heart preparation

The Langendorff technique was used to evaluate the effect of alamandine-(1-5) on left ventricular contractility and coronary arteries. Wistar rats were anesthetized with ketamine (100 mg/kg) and xylazine (10 mg/kg), then decapitated 10–15 min after intraperitoneal injection of 200 IU heparin. Briefly, the thorax was opened and the heart was carefully dissected and perfused through an aortic stump with Krebs-Ringer solution (KRS) containing (in mmol.L^-1^) 118.4 NaCl , 1.2 KH_2_PO_4_, 4.7 KCl, 1.2 MgSO_4_⋅7H_2_O, 1.26 CaCl_2_⋅2H_2_O, 26.5 NaHCO_3_ and 11.7 glucose at 37 °C with constant oxygenation (5 % CO_2_ and 95 % O_2_). A water-filled balloon was inserted into the left ventricle through the left atrium to record the isovolumetric ventricular pressure and the volume was adjusted to keep the left ventricular end-diastolic pressure between 8 and 12 mmHg. The maximal rate of left ventricular pressure rise (dP/dt_max_), maximal rate of left ventricular pressure decline (dP/dt_min_), and heart rate (HR) were calculated from the intraventricular pressure wave. Coronary perfusion pressure was acquired through a pressure transducer connected to the aortic cannula and coupled to a data-acquisition system (DATAQ Instruments, USA). During the stabilization period (20 min), the perfusion flow was adjusted to keep the perfusion pressure between 70 and 100 mmHg. Thereafter, the perfusion flow was maintained constant and the hearts were perfused for an additional 15 min with KRS containing alamandine-(1-5) 20 pmol.L^-1^ or 200 pmol.L^-1^.

#### Cell Culture and transfection

Chinese hamster ovary (CHO) cells were obtained from Cell Bank of Rio de Janeiro (BCRJ) and human umbilical vein cell line (EA.hy926) were kindly donated by Dr. Luciano Capettini (Universidade Federal de Minas Gerais, Brazil). All cells were cultured in DMEM cell media (Thermo-Fisher, Waltham, MA, USA) supplemented with 10% fetal bovine serum (FBS) (Cultilab, Campinas, Brazil) and 1% of antibiotic/antimycotic solution (Thermo-Fisher, Waltham, MA, USA) at 37°C, 5% CO_2_. Stable transfectants were generated by transfecting CHO cells with 500 ng of plasmid DNA using Lipofectamine 2000 (Thermo-Fisher, Waltham, MA, USA) and selecting with 400 μg.ml^-1^ G418 geneticin (Thermo-Fisher, Waltham, MA, USA), according to the instructions of the manufacturer. Mas, AT1 and AT2 expression constructs were generated by subcloning human MasR, AT1R or AT2R in pcDNA3.1 expression vector as described previously^13^. The human MrgD receptor expression construct was obtained from GeneCopoeia^TM^ (Rockville, MD).

#### Imaging of intracellular NO production in EA.hy926, CHO and Mas, MrgD and AT2R transfected CHO cells

We used nitric oxide (NO) measurements in order to test the alamandine-(1-5) activation of the transfected human receptors in CHO cells and EA.hy926 cells. Cells were grown overnight in glass slides and loaded with 10^-6^ mol.L^-1^ 4,5-diaminofluorescein-diacetate (DAF-2DA; Molecular Probes) for 30 minutes. After washing with PBS to remove excess probe, cells were stimulated for 30 min with Ala-(1–5) at concentrations ranging from 10^−6^ to 10^−8^ mol.L^-1^ or the receptor agonist, leaving untreated cells as controls. Cells were then rinsed in PBS, fixed in 4% paraformaldehyde (w/v) solution for 15 min and then mounted in anti-fading mounting media. Images were obtained in a Nikon Eclipse Ti confocal microscope (Nikon, Melville, NY, USA) with λex = 495 nm and λem = 505 nm. Confocal images fluorescence were quantified by ImageJ14.

#### Evaluation of cardiovascular effects induced by intravenous injections of alamandine-(1-5)

One day before the experiments, animals (SHR and Wistar, 12-14 weeks old, n= 6 each) were anesthetized with a mixture of Ketamine (50 mg/kg) and Xylazine (10 mg/kg). Polyethylene cannulas made with PE10 (4 cm for the artery and 3 cm for the vein) polymerized by heat with PE50 (aprox. 17 cm) and filled with 0.9% sterile saline containing heparin (5000UI/ml) were inserted into the abdominal aorta through the femoral artery (BP recordings) and into the femoral vein (drug injections). Catheters were externalized through the subcutaneous to the interscapular region of the animal and fixed with surgical suture thread. Twenty-four hours later, cardiovascular parameters were recorded in conscious animals. Arterial catheter was connected to a pressure transducer and pulsatile arterial pressure (PAP) was continuously recorded by an A/D data acquisition system (MP100; Biopac Systems, Inc., Santa Barbara, CA, USA). Mean arterial pressure (MAP) and heart rate (HR) were simultaneously derived from arterial pulse waves by software (AcqKnowledge5; Biopac Systems). After a period of one hour of stabilization of the cardiovascular parameters, the peptide alamandine-(1-5) was injected intravenously at a dose of 30 ug/kg. The volume of the injection was 0.1 mL/100g. A similar volume of sterile saline was injected in SHR and Wistar rats. After the injection, cardiovascular parameters were accompanied for 6 hours.

#### High resolution echocardiographic measurements

To perform the echocardiographic examination, animals underwent anesthetic induction in an anesthetic chamber through inhalation of a mixture of oxygen and 5% Isoflurane. After induction, the anesthetic plan was maintained with a mixture of oxygen and 3% Isoflurane, in a spontaneous mask breathing. Following, chest was trichotomized and the animals were positioned in the supine position to acquire echocardiographic images with simultaneous acquisition of the electrocardiographic trace, using the Vevo 2100 echocardiograph (FujiFilmVisualSonics Inc., Toronto, Ontario, Canada) and a 13 MHz linear transducer. To evaluate systolic function, an echocardiographic section was performed in the transverse plane of the left ventricle, at the level of the papillary muscles. Using the two-dimensional mode, the fractional area change of the left ventricular cavity was evaluated. Using M mode, the ejection fractions and shortening of the left ventricle were evaluated, as well as the thickness of the anterior wall and posterior wall of the left ventricle in diastole and systole, from which ventricular mass was also calculated. To evaluate diastolic function, an apical 4-chambers, longitudinal section, of the left ventricle was made and the sample was positioned above the mitral valve to evaluate the left ventricular filling flow. The speed of movement of the septal ventricular wall was also assessed using tissue Doppler. To evaluate cardiac output, longitudinal echocardiographic outflow tract was obtained, from which the VTI was calculated. It used the formula: [(DiamLVOT X π) X VTI] X HR for the calculations. Echocardiographic measurements followed the standards of the Standardization Committee of the American Society of Echocardiography.

#### Evaluation of cardiovascular effects induced by ICV infusion of alamandine-(1-5)

Rats (SHR and Wistar, 12-14 weeks old) were anesthetized with a ketamine (50 mg/kg) and xylazine (10 mg/kg) mixture (i.p.) and subjected to stereotaxic implantation of the metallic cannula (25 gauge) into the lateral ventricle and cemented with three anchoring screws to the skull. The tip of the cannula was placed 1.0 mm caudal to the bregma, 1.5 mm lateral to the midline, and 4.5 mm ventral to the dorsal surface of the skull, using the atlas of Paxinos & Watson ^15^ as a reference. ICV cannula was connected to a polyethylene tubing (PE-10), filled with sterile isotonic saline, and fixed to the interscapular region. After surgery, rats received a poly-antibiotic (20U; Pentabiotico®, Fort Dodge, Brazil) and flunixin meglumine (1 mg/Kg, s.c.; Banamine®, Schering Plough, Brazil) for post-operation analgesia. The site of infusion was verified postmortem by the presence of Alcian blue dye (5%), injected through the ICV cannula (2 μL), only in the ventricular system. Cardiovascular parameters recording took place 5 to 7 days after stereotaxic surgery. One day before the experiments, under ketamine (50 mg/kg) and xylazine (10 mg/kg) anesthesia, the animals underwent implantation of polyethylene cannulas in the femoral artery and vein to recording of cardiovascular parameters and baroreflex control of HR test. Twenty-four hours later, a period of one hour of stabilization of the cardiovascular parameters was given and baseline recording of MAP and HR and baroreceptor reflex test were performed. Baroreflex was evaluated by measuring the reflex changes in HR (converted to pulse interval, PI) in response to transient increases in MAP (mmHg) induced by bolus injections of increasing doses of phenylephrine (PHE, 0.1 ml of 1 - 10 µg/ml, i.v.) as described previously ^16^. Baroreflex sensitivity index of each rat was calculated by the average of the ratio (ΔPI/ΔMAP; ms/mmHg) of the maximal changes in PI and MAP with each dose of PHE. ICV infusion (4 μg/12 μL/h, for 90 min) of alamandine-(1-5) or Saline (NaCl 0.9%; control) was performed through a Hamilton® syringe coupled to an infusion pump. Rats were randomly assigned to either an alamandine-(1-5) or a control group. At the end of the ICV infusion, baroreflex sensitivity was assessed again.

#### Microinjection of peptides into insular cortex

The surgeries were done under anesthesia with urethane (1.4 g/kg i.p.). During the protocol the body temperature was kept in the range of 36.5 to 37.0° C with a heating pad (Physitemp TCAT-2DF Controller). The trachea was cannulated to maintain the airways open. Cannulas made with polyethylene tubing (PE10 connected to PE50) were implanted in the femoral artery, to record mean arterial pressure (MAP), and in the femoral vein for drugs injection or supplemental anesthesia when necessary. Blood pressure was recorded using a pressure transducer (MLT 0699, AD Instruments). The electrocardiogram (ECG) was done using the protocol of Sgoifo and colleagues ^17^ for measurement of heart rate (HR). Using a retroperitoneal approach, the left renal nerve was isolated and covered with mineral oil. The nerve was placed on a silver bipolar electrode to record renal sympathetic nerve activity (RSNA). The signal was amplified by 10K, filtered (100-1000 Hz), displayed on an oscilloscope (Tektronix 546B) and monitored by an audio amplifier. The filtered nerve activity signal was rectified, integrated (resetting every second), and displayed. The noise level of the RSNA recording system was determined post mortem and subtracted from RSNA values obtained during the experiment. Signals - ECG, MAP and RSNA - were collected and continuously displayed on a computer connected to Powerlab 4/20 - LabChart 7.3.2 software (ADInstruments, Sydney, Australia). The animals were positioned in a stereotaxic frame, and a small unilateral craniectomy was made to allow the insertion of a glass pipette (Sigma-Aldrich) into the insular cortex (coordinates anteroposterior +1.5mm/ laterolateral 5.4mm/ dorsoventral -7.0mm). After all surgical procedures, a minimum period of 20 minutes was waitedheld for cardiovascular parameter stabilization. The duration of alamandine-(1-5) (400 pmol/100nL) microinjection into the IC was less than 10 seconds. Because alamandine (40 pmol/100 nL, n=7) evoked a cardiovascular response just at rostral IC, all experiments were performed only at this region. We evaluated changes in HR (beats/min), MAP (mmHg), and RSNA (% baseline) evoked by unilateral microinjection of saline (NaCl 0.9%, *n* = 4) and alamandine-(1–5) (400 pmol/100 nL, *n* = 7) into the rostral IC. All experiments were performed in separate groups of animals.

### Mass spectrometry

#### Blood collection

Blood samples from adult mice were collected from male and female animals (n=7 for each gender) by decapitation into tubes containing protease inhibitors (Roche Life Sciences, Basel, Switzerland) and 0.5% EDTA (v/v).

Blood samples from young volunteers from both sex (28±6 years old and 67±15 kg) were collected by venipuncture in vacuum tubes containing EDTA and protease inhibitors at the same day of the clinical assessment (between 8–11 AM). Blood was immediately centrifuged at 3000 rpm for 10 min, 4°C. Plasma samples were obtained and stored at -70°C until assayed.

Five milliliters of peripheral blood samples from healthy and nephropatic pediatric patients were drawn by venipuncture in vacuum tubes containing EDTA at the same day of the clinical assessment (between 8–11 AM). Blood was immediately centrifuged at 3000 rpm for 10 min, 4°C. Plasma samples were obtained and stored at -70°C until assayed. The demographic characteristics of the patients are shown on table 1 (supplementary data).

This study was approved by the Research Ethics Committee of the *Universidade Federal de Minas Gerais*, Brazil (Protocol number: CAAE-0547.0.203.000-11). The legal responsible of all participants (patients and controls) provided written informed consent before admission to the study. The study protocol did not interfere in any medical procedure. Follow-up and treatment were guaranteed even in case of refusal to participate in the study.

#### Sample preparation for angiotensin quantification in plasma

Samples were centrifuged (800xg x at 4°C for 10 minutes) and plasma was stored at −80°C until extraction. Solid Phase Extraction (SPE) was performed using the Sep-Pak C183cc Vac RC Cartridge, 500 mg sorbent (Waters, Milford, MA, USA). The C18 resin was activated by 2 sequential washes with 10 mL of 99% acetonitrile (ACN) / 0.1% formic acid (FA) (v/v), followed by 10 mL of 0.1% FA (v/v). Then, the resin was conditioned with 3 mL of 0.1% FA / 0.1% BSA (v/w), 10 mL of 10% acetonitrile / 0.1% FA (v/v), and 3 mL of 0.1% FA (v/v). The samples were then loaded and washed with 20 mL of 0.1% FA (v/v). For elution, 3 mL of 60% ACN / 0.1% FA (v/v) was loaded and collected into low-binding polypropylene tubes (Eppendorf, Hamburg, Germany). Samples were dried out using a SpeedVac (Eppendorf, Hamburg, Germany). Dried samples were resuspended in 100 µL 0.1% formic acid (v/v) (concentration factor of the sample = 10-fold).

#### Multiple Reaction Monitoring (MRM) analysis

Chromatography was performed on an ACQUITY I-Class UPLC system (Waters, Milford, MA, USA) coupled to electrospray ionization (ESI) tandem mass spectrometry (LC-ESI-MS/MS Xevo TQ-S (Waters, Milford, MA, USA). The chromatographic separation was done in a C18 column (ACQUITY UPLC BEH C18 Column, 130 Å, 1.7 μm, 2.1 mm X 100 mm, Waters, Milford, MA, USA) for 5.5 min, each injection (10 µL volume per sample). Solvent A was made of 0.1% formic acid in H_2_O (v/v) and solvent B of 0.1% formic acid in ACN (v/v). The chromatographic gradient was set as previously described ^18^ in a 300 µL/min flow rate as follows (expressed as % of solvent B): i) 3-40% in 3.5 min, ii) 40-99% in 0.01 min, iii) 99% for 0.99 min, iv) 99-3% in 0.01 min, and v) 3% for 0.99 min. Regarding the MS analysis, the main parameters were as follows: i) capillary = 3.5 kV; ii) cone = 20V; iii) temperature of the desolvation gas (hydrogen) = 550 °C. The collision energy (CE) (argon gas) was tuned for each target peptide spanning from 10 to 20 CE. The following transitions were monitored by the MS in the MRM mode. Alamandine: 286.1 <136.1 and 286.1 < 327.2; Ang A: 335.1 < 231.8 and 335.1 < 513.2; alamandine-(1–5): 621 < 210.94 and 621 < 508.1 m/z. The calibration curve was obtained using a stock solution containing a mixture of synthetic peptides. The applied calibration curve model (y = ax + b) proved accurate over the concentration range 10 to 1000 pg·mL^−1^(r=0.997). Data acquisition was performed using the software MassLynx MS (Waters, Milford, MA, USA) and raw data evaluation by its application TargetLynx™.

#### MALDI TOF/TOF analysis

In order to see whether alamandine is metabolized from ECA forming alamandine-(1-5), we took advantage of the availability of the global angiotensinogen deficient mice (TLM) ^19^. Blood samples were collected no from TLM mice in heparin tubes and centrifuged (5000 RPM, 10 min, 4°C) for plasma separation. In tubes containing 470 µL of Tris-HCl buffer pH =7.4, we added 20 µL of TLM mice plasma and 10 µL of alamandine solution at 500 µg.mL^-1^ (final concentration 10 µg.mL^-1^). Samples were incubated at 37°C for 0, 1 and 3 minutes, with or without 1.2 μmol.L^-1^ of the ACE inhibitor, Captopril (Signa-Aldrich). Reaction was stopped by addition of a protease inhibitor cocktail (Roche Life Sciences, Basel, Switzerland) and EDTA 0.5% (v/w) immediately at due time. Samples lacking alamandine solution were used as controls. Prior to mass spectrometry analysis in an Autoflex III Smartbeam MALDI TOF/TOF spectrometer (Bruker Daltonics, Billerica, USA), we proceed to sample desalting in C18 P10 ZipTip® Pipette Tips (Millipore, Billerica, MA) following manufacturer instructions. Samples were eluted directly in a MTP 384 Polished Steel target plate (Bruker Daltonics, Billerica, USA) and diluted (1:1, v/v) with α-cyano-4-hydroxycinnamic acid matrix (Sigma-Aldrich, Saint Louis, USA). Sample-matrix co-crystallization was carried out by the dried-droplet method ^20^. External calibration was performed by homogenizing the Peptide Calibration Standard II (Bruker Daltonics, Billerica, USA) in 1:1 ratio in α-Cyano-4-hydroxycinnamic acid matrix directly on the same target plate. MS spectrum was obtained in positive reflected mode with 400 laser shots and an acceleration voltage held at 9.71–21.01 kV.

#### Statistical analysis

The statistical analysis for each method is described in the figures legends, including one-way ANOVA, two-way ANOVA, paired and non paired *t*-test and nonparametric test. The criterion for statistical significance was set at p value < 0.05.

## RESULTS

### Opposing effects of alamandine and alamandine-(1-5) in mouse aortic rings

Considering that alamandine produces vasorelaxation of mouse aortic rings we took advantage of angiotensin receptors KO in our laboratory to start the alamandine-(1-5) evaluation using mouse aortic rings. Strikingly, the pentapeptide produced a significant increase in the aortic rings tension (Fig. 1A). The contraction was already evident at a concentration of 10^-11^ mol.L^-1^. The contractile effect was only slightly increased in endothelium-denuded rings (Fig. 1B). Contrasting with aortic rings, alamandine-(1-5) produced a concentration-dependent relaxation in the mouse small mesenteric artery (Fig. 1C-D).

**Figure 1.**
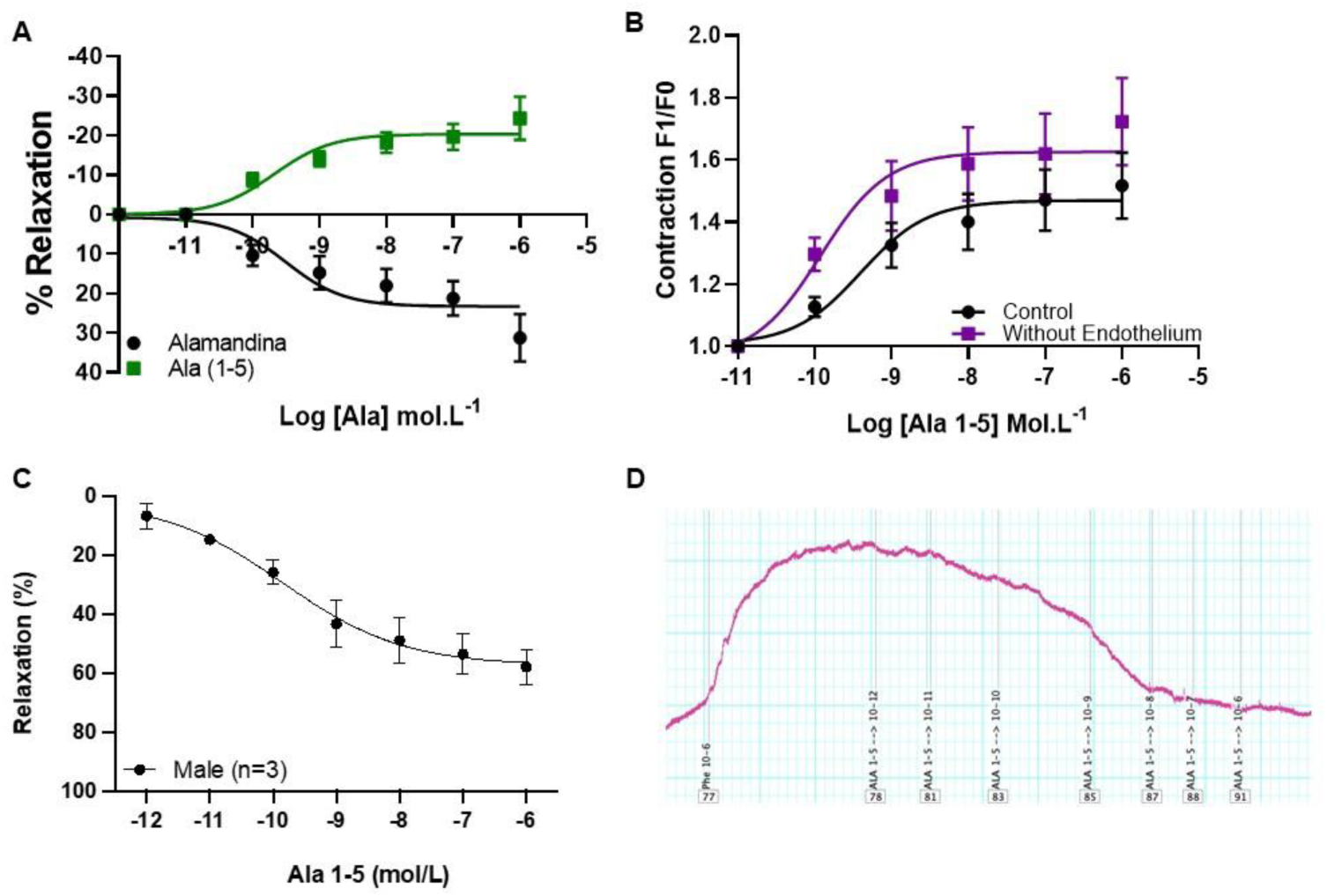
Vascular effects of Alamandine-(1-5) in isolated vessels. A. Cumulative concentration of Ala or Ala-(1-5) in aortic rings; B. Effects of Ala-(1-5) with intact and denuded endothelium in aortic rings; C. Effects of Ala (1-5) in microvessels. D. Representative image of recording vascular reactivity in response to Ala-(1-5) in a microvessel. Data were expressed as mean ± SEM and analyzed by two-way anova followed by Fisher’s multiple comparisons test. p<0.05.

### Alamandine-(1-5) induces a dramatic decrease in ventricular myocyte contractility

Given the instigating results reported above, we next assessed the effect of alamandine-(1-5) in isolated ventricular myocytes stimulated to contract at 1Hz. Cells were incubated with a range of alamandine-(1-5) starting from 10 nmol.L^-1^ until 1 μmol.L^-1^ for 10 min. As shown in Fig. 2A, alamandine-(1-5) induced a concentration-dependent decrease in the shortening area of ventricular myocytes, with more pronounced effects observed in cells exposed to 100 nmol.L^-1^ (Fig. 2 A). With this in mind, we opted to use for the next series of experiments a concentration of alamandine-(1-5) of 100 nmol.L^-1^. Samples motion vector images acquired from cardiomyocytes exposed or not to alamandine-(1-5) 100 nmol.L^-1^ are presented in Fig. 2B-C. As shown in Fig. 2D-F alamandine-(1-5) induced a significant decrease in shortening area and speed parameters. To assess whether the reduction in contractility is due to alterations in Ca^2+^ release, we next recorded Ca^2+^ transient amplitude by confocal microscopy in cells loaded with Fluo-4 AM. Consistent with the contractility data, cardiomyocytes exposed to alamandine-(1–5) presented reduced and slower Ca^2+^ transients. Taken together, the compelling in vitro data obtained so far with alamandine-(1–5) indicated that this peptide exerts unique actions that are distinct from alamandine, suggesting the idea that alamandine-(1–5) is a new biologically active component of the RAS.

**Figure 2.**
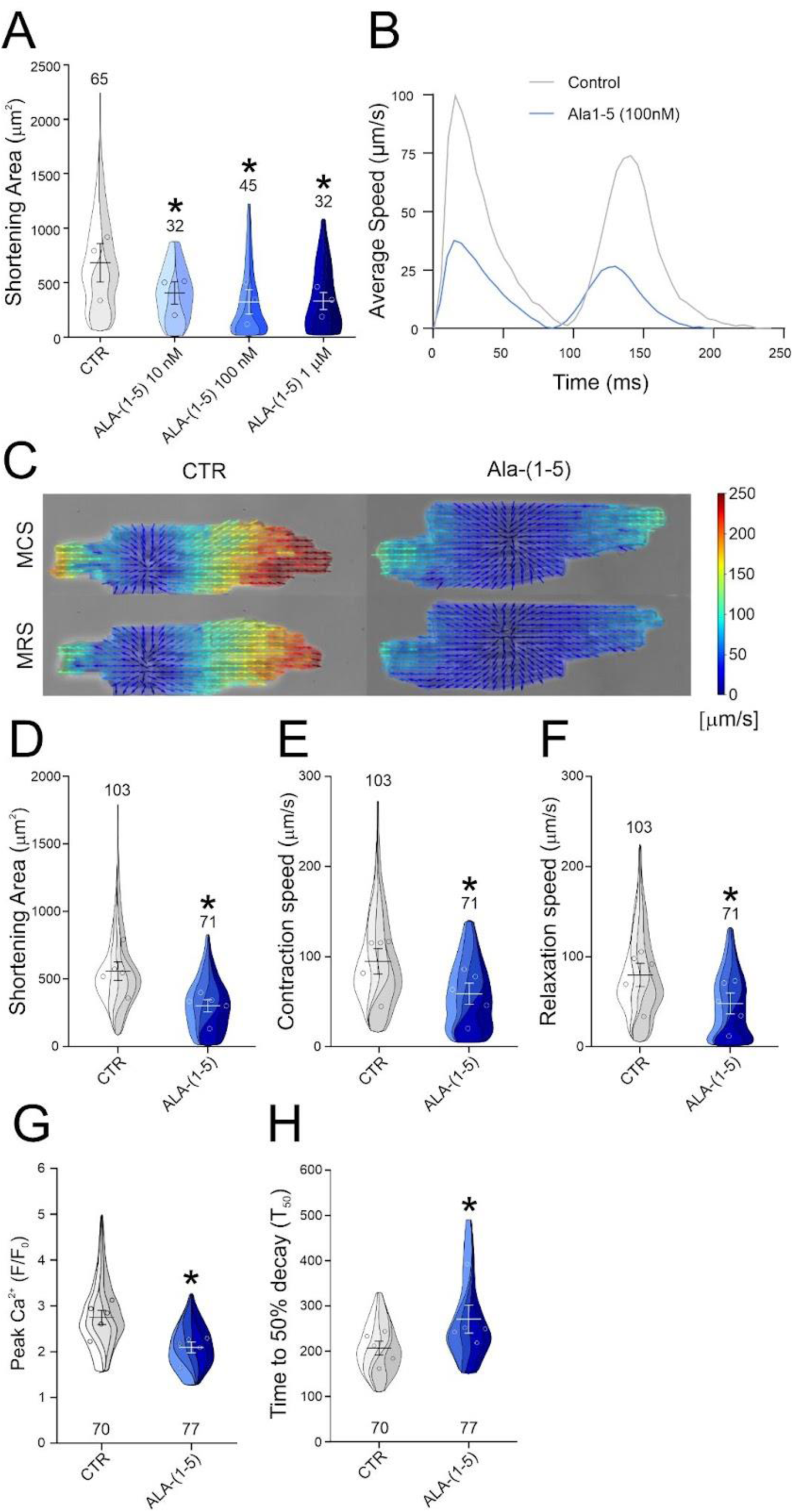
Alamandine-(1-5) reduces ventricular myocyte shortening area. A. Concentration-dependent effect of alamandine-(1-5) in isolated ventricular shortening. B. Sample average-speed trace obtained from a single contraction-relaxation cycle from control and under the effect of alamandine-(1-5). C. Cardiomyocyte displacement speed displayed in magnitude and vector fields with visual and numerical intensity scale detection obtained during maximum contraction speed (MCS) and maximum relaxation speed (MRS) in response to alamandine-(1-5). D-F. A marked decrease in cardiomyocyte shortening and speed parameters was observed in response to alamandine-(1-5) treatment. G-H. Alamandine-(1-5) induced a decrease in Ca^2+^ transient amplitude while it increased the kinetics of decay of the Ca^2+^ transient (T50). In violin plots solid lines and dashed lines indicate medians and 25th and 75th percentiles, respectively. Colored shapes show density estimates of the data. In A *p<0.05 In D-H, * p<0.05 Student’s *t*-test when compared to untreated control.

### Alamandine-(1–5) reduced heart contractility in rat Isolated hearts

Given the data obtained with isolated myocytes and aiming to get further insight on the effect of alamandine-(1-5) in the heart contractility, we next used the Langendorff technique. As shown in Fig. 3A, alamandine gradually reduced LEVSP at a dose of 20 pmol.L^-1^. This effect was transient at a dose of 200 pmol.L^-1^ (Fig 3A). LEVDP was increased by alamandine-(1-5) 200 pmol.L^-1^. No significant effects in the LVEDP were observed at the dose of 20 pmol.L^-1^ (Fig. 3B). Interestingly, the pentapeptide at the dose of 200 pmol.L^-1^ evoked a biphasic effect on dP/dt_max_ (Fig. 3C) and dP/dt_min_ (Fig 3D), i.e., a rapid and transient increase followed by a significant decrease in these parameters. At a dose of 20 pmol.L^-1^, alamandine-(1–5) only produced a gradual reduction in these parameters. Alamandine-(1-5) 200 pmol.L^-1^ induced a reduction in perfusion pressure at the beginning of the perfusion period, indicating coronary vasodilation, which was returned to baseline values at the end of the analyzed period (Fig. 3E). In contrast, with the dose of 20 pmol.L^1^, alamandine-(1-5) promoted an initial slight, but not significant, reduction in the perfusion pressure followed by a progressive increase in this parameter. No effects were observed on HR (Fig. 3F).

**Figure 3.**
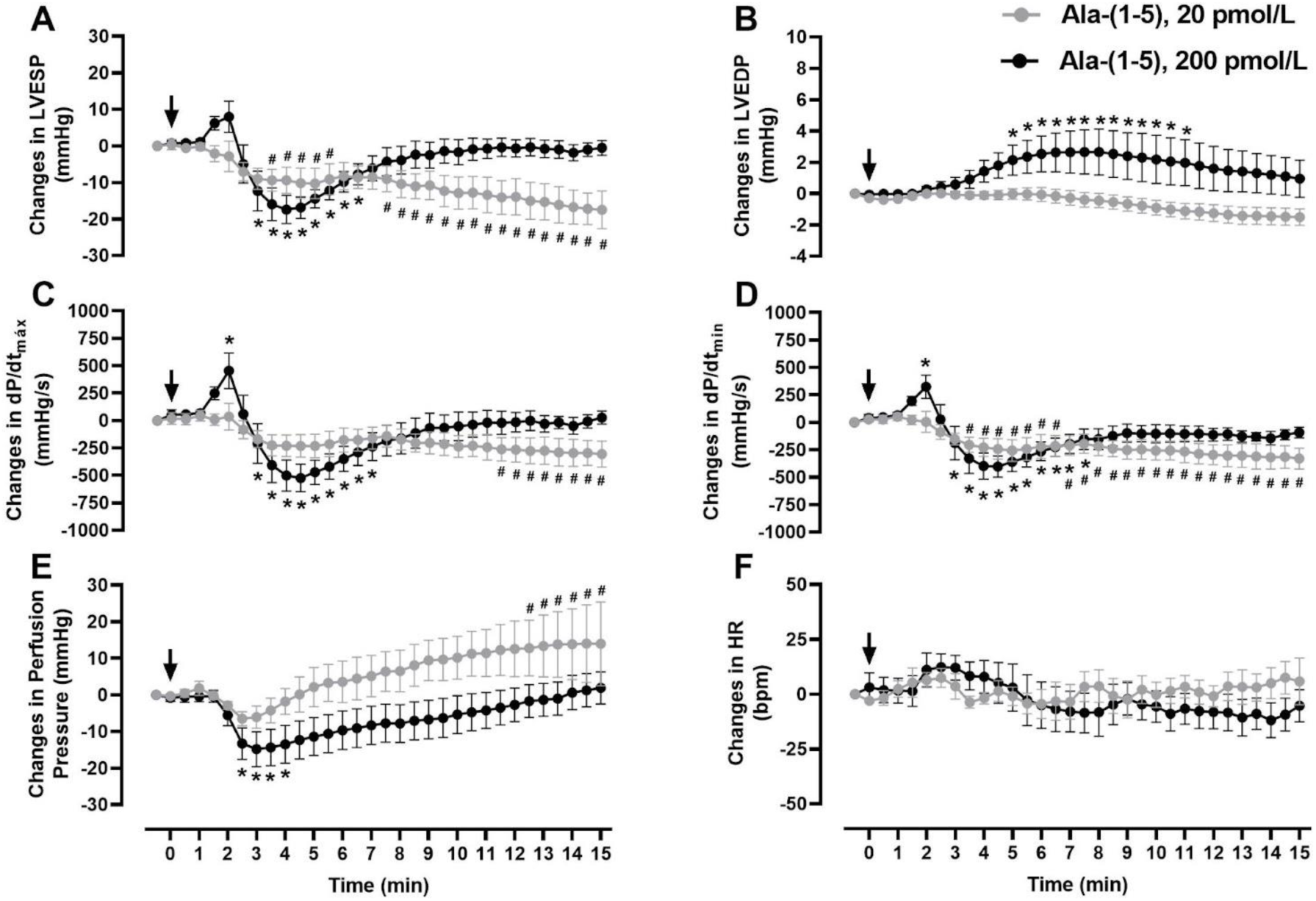
Effects of alamandine-(1-5) in isolated rat hearts. After a stabilization period (20-30 min) the hearts were continuously perfused with alamandine-(1-5) at doses of 20 or 200 pmol/L for 15 min. The graphs show the changes in (A) left ventricular end-systolic pressure (LVESP), (B) left ventricular end-diastolic pressure (LVEDP), (C) maximal rate of left ventricular pressure rise (dP/dt_max_), (D) maximal rate of left ventricular pressure decline (dP/dtmin), (E) perfusion pressure, (F) heart rate (HR). Arrows indicate the beginning of the perfusion with alamandine-(1-5). #p<0.05 (20 pmol/L) and *p<0.05 (200 pmol/L) compared with the baseline of the respective group. Statistical significance was calculated by two-way ANOVA followed by Dunnett’s multiple comparisons test. Each point represents the mean ± S.E.M.

### Central effects

#### Cardiovascular and reflex effects of acute ICV infusion of alamandine-(1-5)

In conscious freely-moving rats, 90 minutes of alamandine-(1–5) infusion did not significantly alter baseline values of MAP or HR in normotensive (MAP: 103±8 mmHg, before and 105±11 mmHg, 90 min infusion; HR: 312±13 bpm, before and 331±10 bpm, 90 min infusion) or in hypertensive rats (MAP: 159±8 mmHg, before and 161±10 mmHg, 90 min infusion; HR: 296±7 bpm, before and 310±2 bpm, 90 min infusion). As expected, SHR animals presented a marked attenuation of the baroreflex control of HR (0.3±0.1 ms/mmHg) compared with Wistar rats (1.5±0.5 ms/mmHg). As shown in Fig. 4, alamandine-(1-5)-infused SHR animals presented a significant improvement in the sensitivity of the baroreflex bradycardia (0.5±0.1 ms/mmHg) compared with the same animals before alamandine-(1-5) infusion (0.3±0.1 ms/mmHg). In contrast, alamandine-(1-5) ICV infusion did not significantly change baroreflex control of HR of Wistar rats (1.4±0.3 ms/mmHg). Cardiac autonomic balance was evaluated by means of the pharmacological blockade of cardiac muscarinic and 𝛽-adrenergic receptors. Interestingly, alamandine-(1-5) did not produce significant changes in cardiac autonomic balance both in normotensive and hypertensive strains (Fig. S1, Supplementary Data).

**Figure 4.**
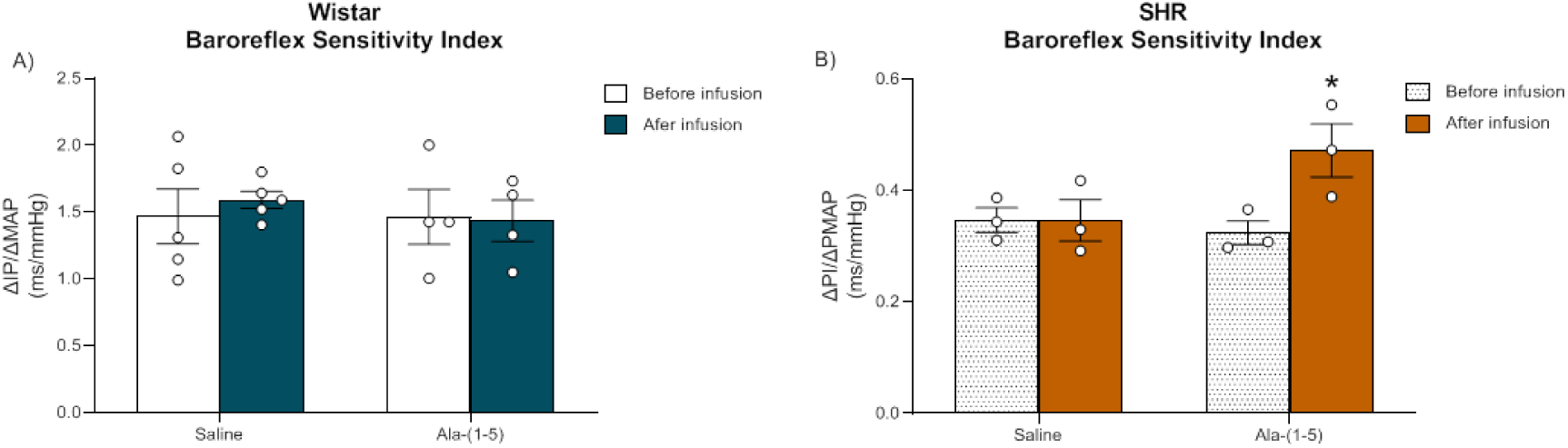
Alamandine-(1-5) improves baroreflex in SHR. Index of the sensitivity of baroreflex bradycardia (ΔPI/ΔMAP, ms/mmHg) calculated by the ratio between baroreflex-mediated changes in heart rate (converted to pulse interval, in ms) by changes in mean arterial pressure (ΔMAP, in mmHg) produced by bolus intravenous injections of phenylephrine (1, 2.5, 5, 10 and 20 μg/ml), before and 90 minutes the ICV infusion (12 µl/hour) of saline or alamandine-(1-5) [Ala-(1-5), 0.04 µg/µl] in Wistar (A) and SHR rats (B). Values are mean ± S.E.M. * p>0.05 *vs*. before infusion (paired *t*-test).

#### Effects of alamandine-(1-5) in the Insular Cortex

We have recently shown that the insular cortex is a brain site were alamandine has unique effects (increase in blood pressure, heart rate and renal nerve activity) contrasting with the essentially absent effects of Ang-(1-7). Thus, we next aimed to determine the effect of alamandine-(1-5) in this region. As shown in figure 5, microinjection of this pentapeptide in the rostral part of the insula (AP+1.5mm, from bregma). At the baseline, there was no difference between the groups (vehicle: 85 ± 2 mmHg and 388 ± 5 beats/min, alamandine: 83 ± 9 mmHg and 389 ± 10 beats/min or Ala-(1–5): 91 ± 6 mmHg and 386 ± 7 beats/min). Vehicle (saline, n=4) microinjection was performed as a control for nonspecific autonomic and cardiovascular effects produced by volume microinjection and/or tissue damage. Saline microinjection did not evoke changes in the evaluated parameters (ΔMAP: 2 ± 1 mmHg, ΔHR: 4 ± 2 beats/min, ΔRSNA: 3 ± 1%, Fig. 5). Alamandine (n=7) microinjection into the rostral IC produced a long-lasting (18 ± 3 min) increase in MAP (Δ= 14 ± 2 mmHg), and this effect was associated with a significant increase in HR (Δ= 32 ± 4 beats/min) and in RSNA (Δ= 36 ± 6%). These responses were elicited immediately after alamandine microinjection. When compared with vehicle and, very different from what was observed with alamandine, microinjection of alamandine-(1–5) (n=7) evoked a significant increase in blood pressure without altering renal nerve activity or heart rate (ΔMAP: 22 ± 2 mmHg, ΔHR: -1 ± 3 beats/min, ΔRSNA: 1 ± 3%; Fig. 5). Differently from alamandine evoked responses, the effects of alamandine-(1–5) had a shorter duration (5.88 ± 1.9 min) without delay to beginning the effects.

**Figure 5.**
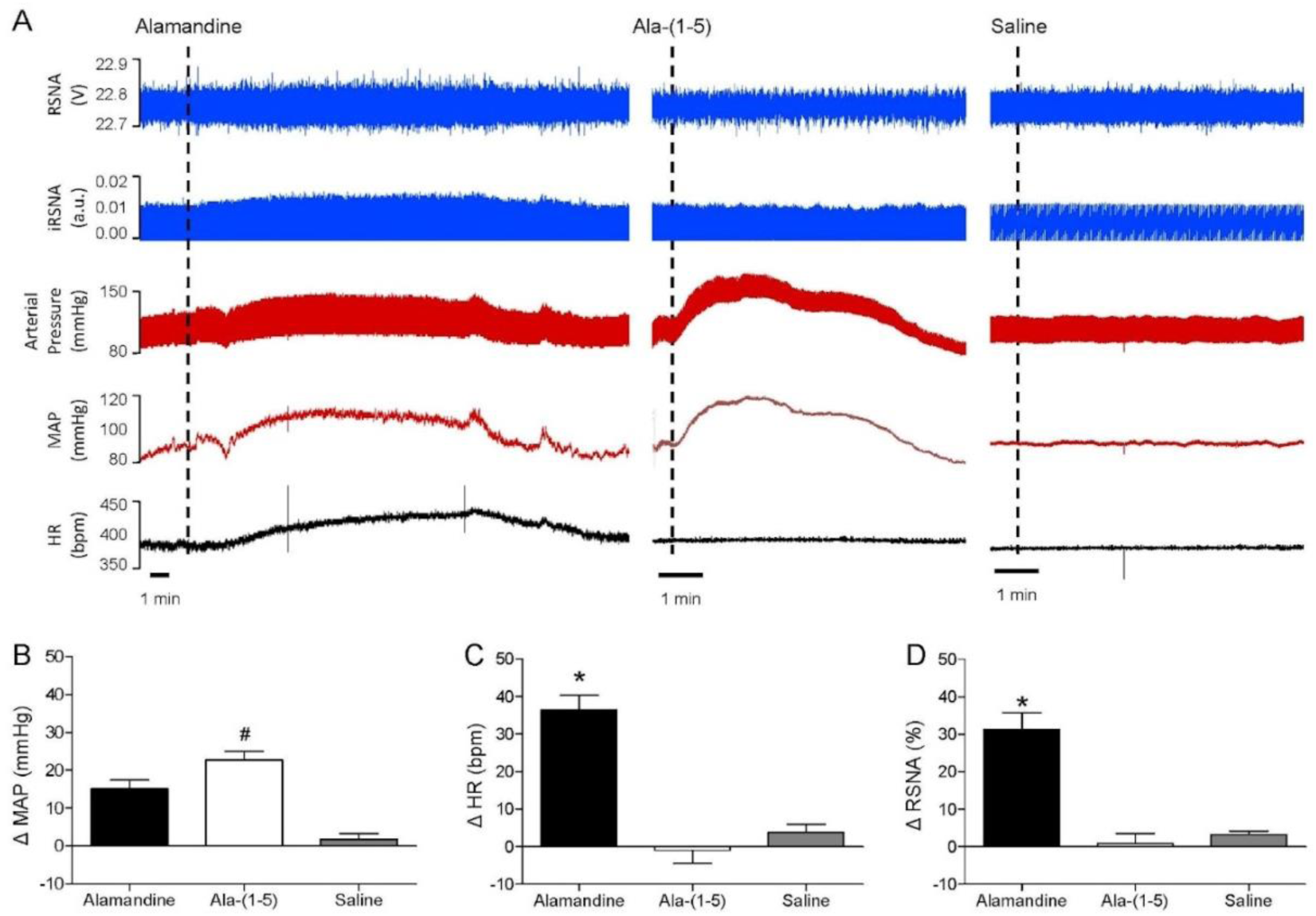
Effect of alamandine-(1-5) in the insular cortex. Representative tracings showing effects on baseline cardiovascular and autonomic parameters (RSNA, renal sympathetic nerve activity, %; iRSNA, integral renal sympathetic nerve activity peak, arbitrary units; arterial pressure, mmHg; MAP, mean arterial pressure, mmHg; HR, heart rate, beats/min) elicited by alamandine (n = 7), alamandine-(1-5) (n=7) and saline (n=4) microinjection into the rostral level of the posterior IC (A). (B): changes in mean arterial pressure (MAP),(C) heart rate (HR) (D), and renal sympathetic nerve activity (RSNA) elicited by alamandine (black bars), alamandine-(1-5) (white bars) and saline (grey bars) microinjection into the rostral level of the posterior IC, coordinates +1.5 mm at rostrocaudal level. Statistical analysis: Anova one way, p < 0.05, * vs. alamandine-(1-5) and Saline # vs. Alamandine and Saline. IC, insular cortex.

#### *In vivo* cardiovascular effects of alamandine-(1-5) in SHR

Based on previous observations showing an anti-hypertensive effect of alamandine in SHR ^2^ we next tested the effect of bolus intravenous (iv) injection of alamandine-(1-5) in SHR and Wistar control rats. As shown in Fig. 6, iv administration of alamandine-(1-5) produced a progressive decrease in systolic blood pressure in freely moving SHR, reaching -33.3±3.0 mmHg, 6 hours post administration (systolic blood pressure: 208±7.6 mmHg, before and 175±10.5 mmHg, 6 hours post injection). This marked effect in blood pressure was not accompanied by significant changes in HR (baseline: 348+24.2 beats/min). Alamandine-(1-5) administration in Wistar rats produced only slight changes in blood pressure (99±11.5 mmHg, 6 hours post-injection vs 109±10.1 mmHg, before) without significant changes in HR (303±12.3 beats/min, 6 hours post-injection vs 328±20.5 beats/min, before). Saline injection did not change blood pressure or HR in Wistar or SHR.

**Figure 6.**
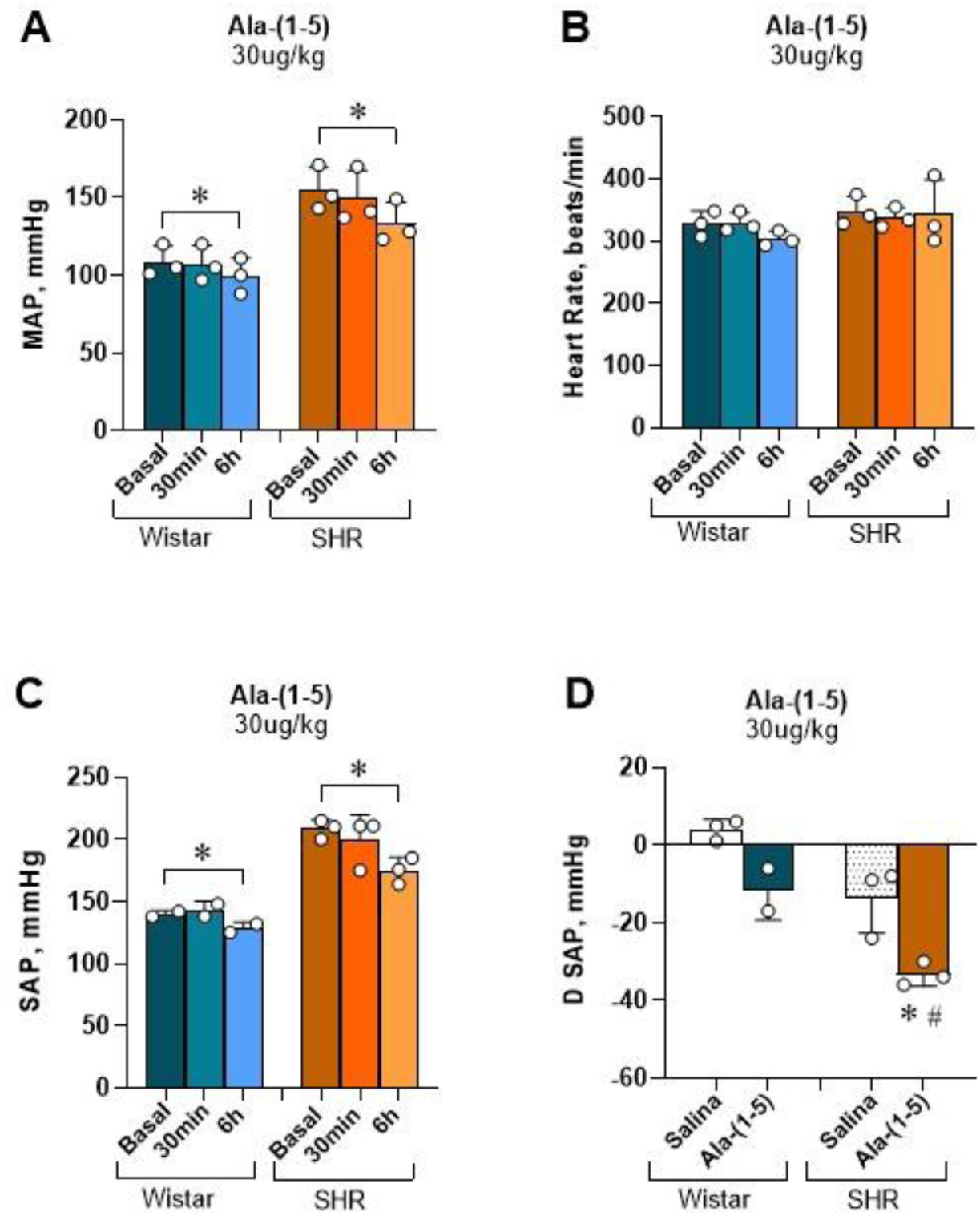
Alamandine-(1-5) anti-hypertensive effect in SHR. Averaged Mean Arterial Pressure (MAP, mmHg; panel A), Heart Rate (HR, beats/min; panel B), Systolic Arterial Pressure (SAP, mmHg; panel C) before (basal), 30 min and 6h after Ala-(1-5) (30 ug/kg) iv bolus injection in Wistar and SHR. Changes in SAP (mmHg; panel D) 6h after iv injection in relation to basal values of Saline and Ala-(1-5) in Wistar or SHR groups. Male Wistar and SHR (n=3 each). Values are mean ± SEM. *p<0.05 vs basal (panels A and C) or Saline (panel D) and #p< 0.05 vs Ala-(1-5) injection in Wistar rats (Two-way ANOVA, followed by Tukey’s post hoc test).

#### High resolution echocardiography in SHR treated with alamandine-(1-5)

In keeping with the results obtained *in vitro* (isolated adult cardiomyocytes) and *ex-vivo* (isolated perfused heart), administration of alamandine-(1-5) in SHR, reduced cardiac output in 33% (from 109.9 ± 10,40 to 74.1 ± 6.02 mL.min^-1^) due to a reduction in 30% of stroke volume (from 279.9 ± 20.98 to 194.6 ± 13.85 uL) two hours after administration. We also observed a reduction in the left ventricle posterior wall and in the E’ wave (Fig. 7F and 7I), respectively.

**Figure 7.**
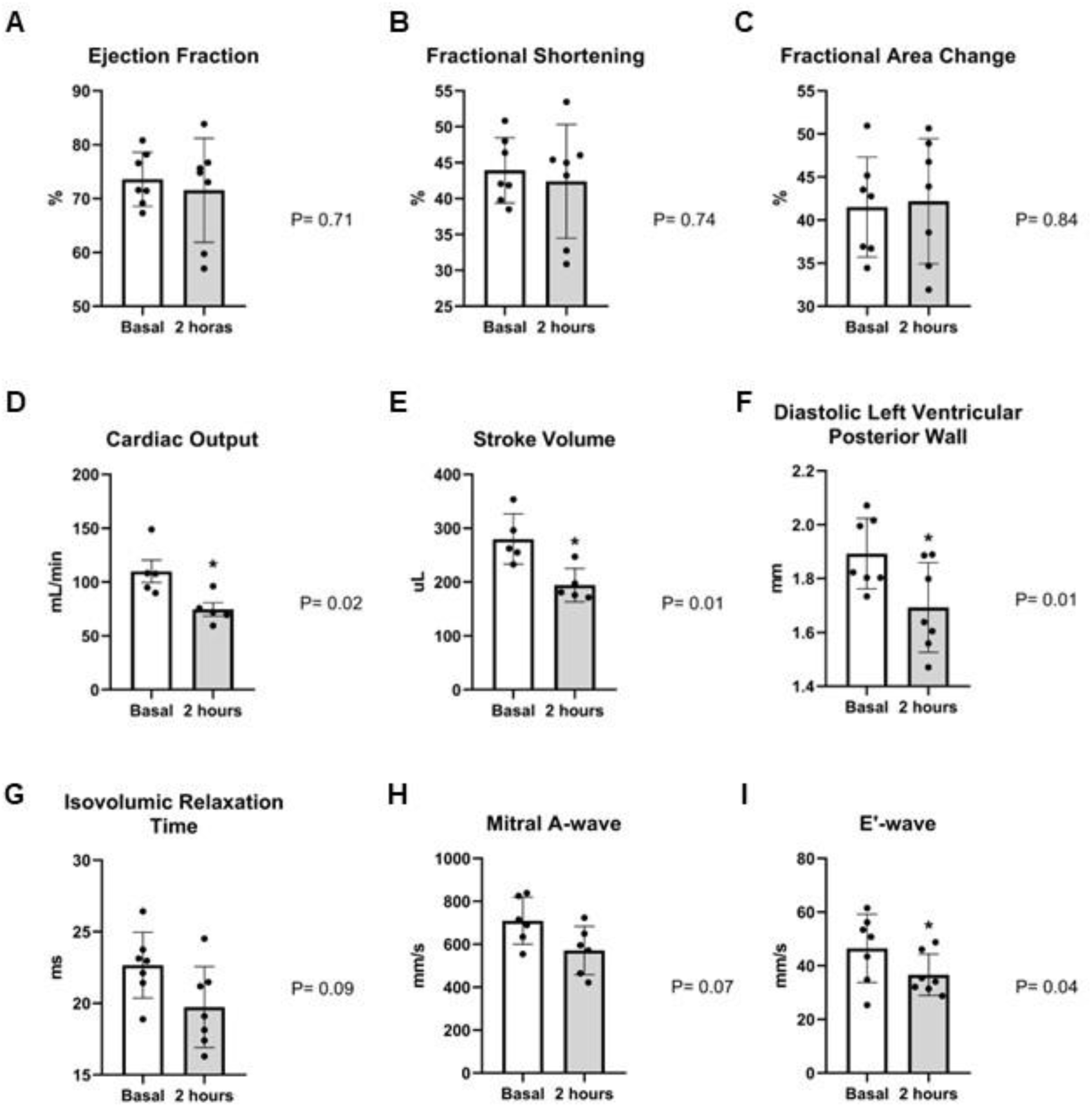
High resolution echocardiography in alamandine-(1-5)-treated SHR. Comparison between baseline and post administration were performed using t-test.

#### Evaluation of the mechanism of action of alamandine-(1-5)

We next used pharmacological and genetic tools to get insights into the mechanism of action of alamandine-(1-5) using mouse aortic rings and cell culture. As shown in Fig. 8A, pre-treatment of aortic rings with L-NAME only slightly increased the contractile effect of alamandine-(1-5) (Fig. S2A, Supplementary Data). The contractile effect of alamandine-(1-5) was essentially abolished in aortic rings taken from MrgD-deficient mice and markedly attenuated in aortic rings taken from AT2-KO mice. In aortic rings from Mas-KO mice the contractile effect of alamandine-(1-5) was not significantly changed (Fig. 8). Likewise, pre-treatment with losartan did not alter the contractile effect of alamandine-(1-5) in this preparation (Fig. S2B, Supplementary Data).

**Figure 8.**
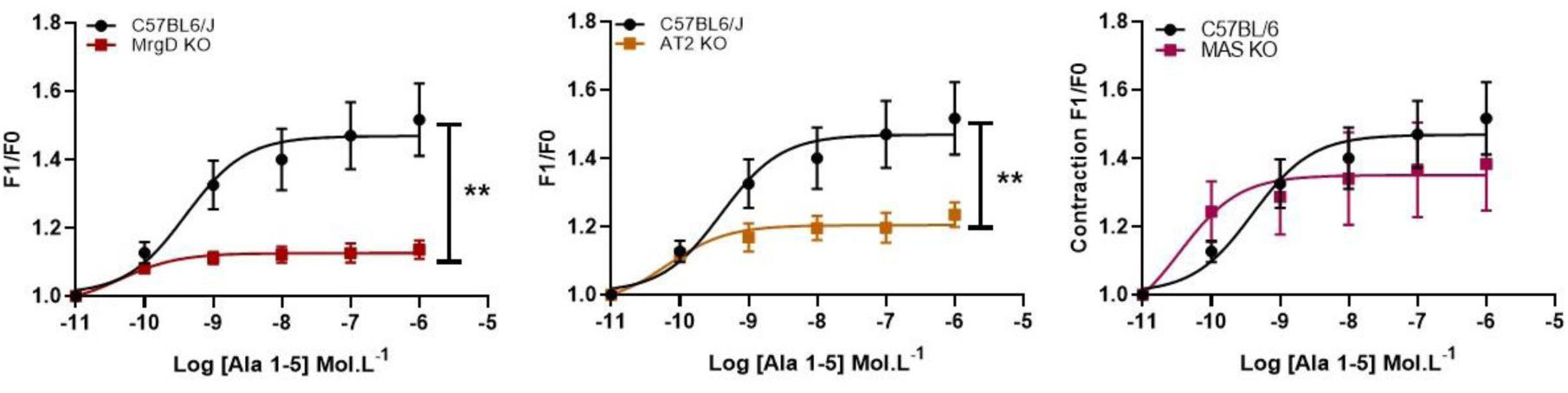
Comparison of the involvement of RAS protective arm receptors in the Alamandine (1-5) response in aortic rings. Effects of ALA (1-5) in aortic rings of MrgD KO (A), AT2 KO (B); and MAS KO (C). Data were expressed as mean ± SEM and analyzed by two-way anova followed by Fisher’s multiple comparisons test. p<0.05.

In order to investigate by which of the classic receptors from renin-angiotensin system alamandine-(1-5) is acting, we used EA.hy926, an endothelial cell line, and CHO cells transfected with either Mas, MrgD or AT2 receptors, to evaluate nitric oxide production upon stimulation with the peptide in comparison with the control unstimulated cells (Fig. 9A-B). NO production was activated by alamandine-(1-5) in EA.hy926 cells by 37% above unstimulated control (p<0.0001). Alamandine-(1-5) also activated CHO cells transfected with Mas, MrgD and AT2 receptors, increasing the NO production by 77%, 85% and 67%, respectively (p<0.0001). Together, this data show that alamandine-(1-5) can activate Mas, MrgD and AT2 receptors.

**Figure 9.**
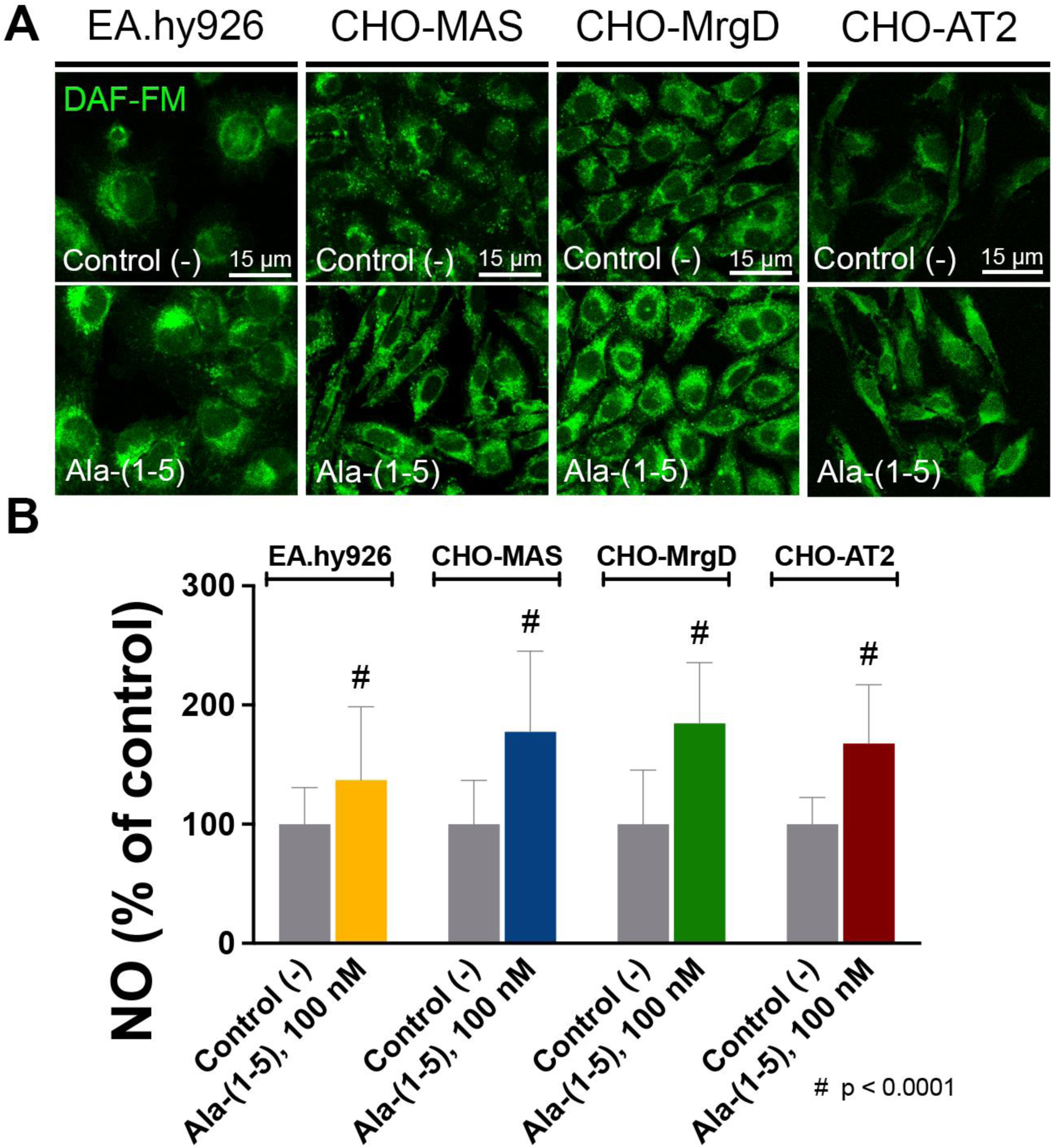
Alamandine-(1-5) activates Mas, MrgD and AT2 receptors. A) Representative images of nitric oxide production in EA.hy926, CHO-MAS, CHO-MrgD and CHO-AT2 cells loaded with DAF-FM and stimulated with alamandine-(1-5) in comparison to the unstimulated control. B) Alamandine-(1-5) can significantly increase NO production in all cell types, indicating that it binds and activates Mas, MrgD and AT2 receptors (p<0.0001; n = 3; at least 90 cells per each condition).

#### Evidence that ACE plays a major role in the formation of alamandine-(1-5) from alamandine in plasma

To explore the potential involvement of ACE in the conversion of alamandine to alamandine-(1-5), plasma samples were collected from Angiotensinogen knockout mice (TLM), which lack angiotensins production. We conducted incubation experiments with alamandine using TLM plasma in presence and absence of the ACE inhibitor captopril. Mass spectrometry analysis (MALDI-TOF/TOF) was then employed to detect any potential formation of a fragment matching the mass of alamandine-(1-5) at 621 Da. As expected, plasma from TLM (TLM_CT) did not present a peak corresponding to angiotensin mass. However, after 1 min (TLM_T1’) and 3 min (TLM_T3’) of TLM plasma incubated with alamandine formation of a peak 621 Da, was observed, corresponding to alamandine-(1-5) mass. In contrast, in the presence of captopril (TLM_T1’_CAP and TLM_T3’_CAP) no formation of a 621 Da peak was detected, highlighting the critical role of ACE activity in the formation of this pentapeptide (Fig. 10A). The same mass spectra was zoomed to the range of 610-650 m/z in order to better visualize 621 Da peak formation in the groups (Fig. 10B). To confirm the identity of spiked alamandine, the pure synthetic peptide and the synthetic peptide in the plasma was submitted to MS/MS fragmentation and sequenced using the *de novo* sequencing method (Fig. 10C). To confirm the identity of alamandine-(1-5) the pure synthetic peptide and the 621 Da peak in the plasma was submitted to MS/MS fragmentation and sequencing (Fig. 10D). Altogether, these results suggest that ACE is responsible for rapidly forming alamandine-(1-5) from alamandine.

**Figure 10.**
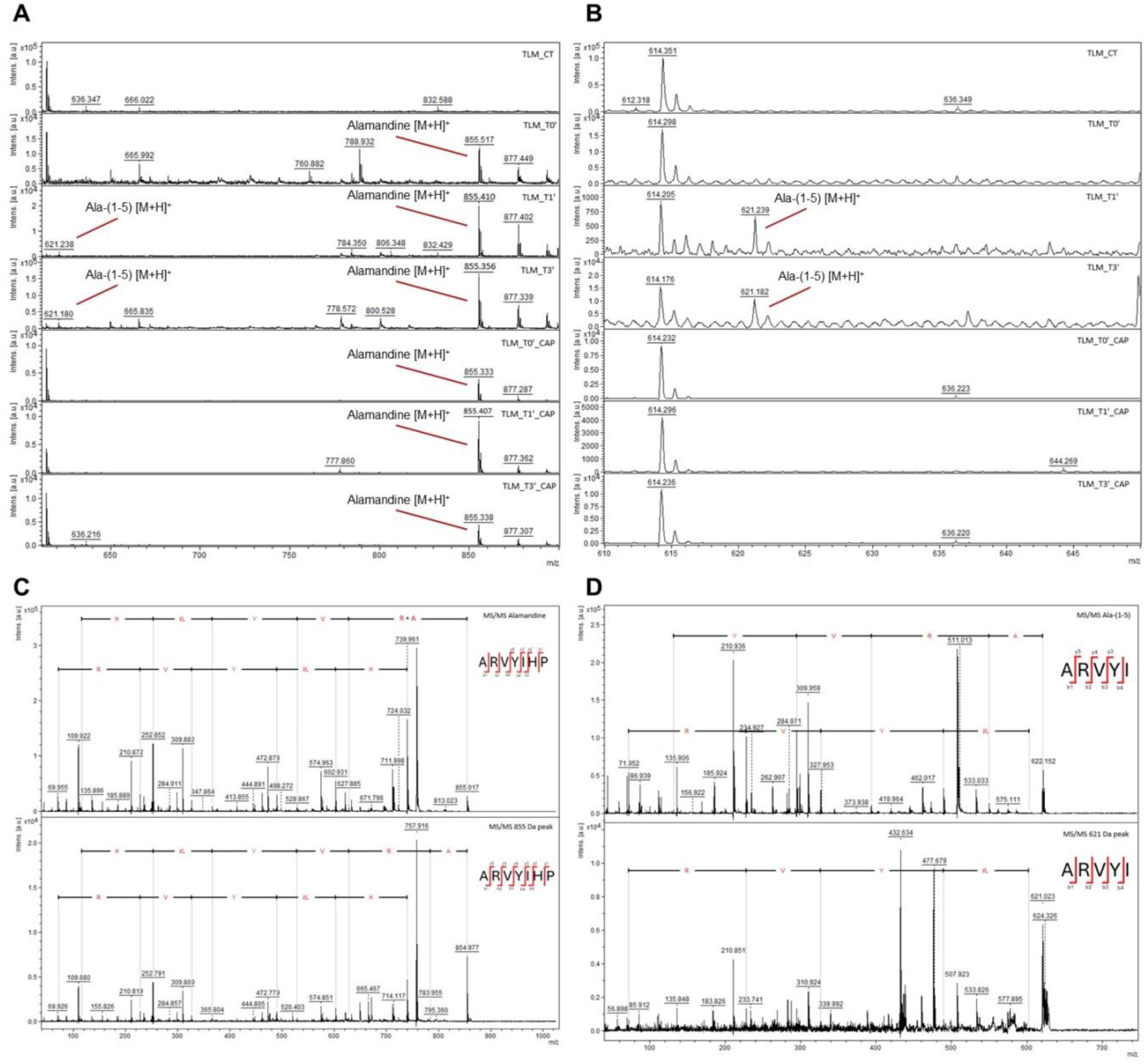
Evidence that ACE is able to metabolize alamandine forming alamandine-(1-5). A. Plasma MS spectra range 610-900 m/z. B. Plasma MS spectra range 610-650 m/z. C. *De novo* sequencing of synthesized alamandine (upper panel) in aqueous solution and spiked alamandine in plasma (bottom panel). D. *De novo* sequencing synthesized alamandine-(1-5) (upper panel) in aqueous solution and formed alamandine-(1-5) in plasma (bottom panel). Alamandine-(1-5) formation by plasma from Angiotensinogen knockout mice (TLM mice) from alamandine was suppressed in presence of the ACE-inhibitor captopril (Panel A and B).

#### Alamandine circulates in mouse and human blood

We next used LC-MS/MS to evaluate whether alamandine-(1-5) is present in human and mouse plasma samples. As shown in Fig. 11A-B we were able to detect alamandine-(1-5) in blood samples from healthy human volunteers and from C57BL6/J mice. The peptide was present in low concentrations in humans ranging from 0.2 to 8.7 pg/mL (average 2.4 ± 1.3 pg/mL). In those samples, the concentration of alamandine-(1-5) precursor, alamandine, ranged from 0.1 to 18.6 pg/mL (average 3.7 ± 2.5 pg/mL). Alamandine-(1-5) was also detected in extracted mouse plasma samples (Fig. 11B).

**Figure 11.**
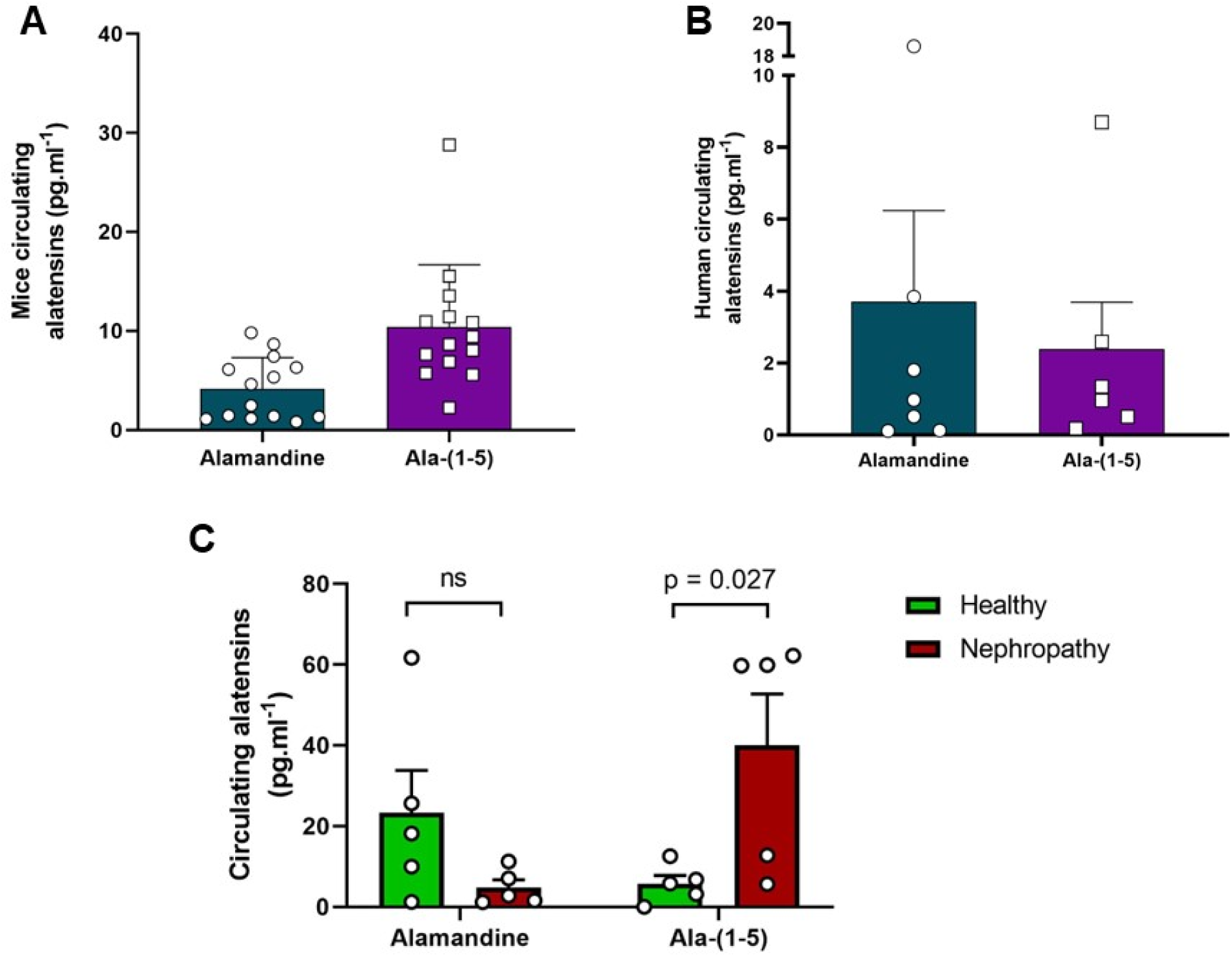
Alamandine-(1-5) is circulation in mice and human plasma. A. Alamandine and alamandine-(1-5) quantification in mice plasma. B. and in humans. Average ± SE of circulating alatensins in healthy subjects. Data obtained by UPLC-MS/MS peptide analyses as previously described ^18^. For comparison between males and females mice t-test was used. **** = p < 0.0001.

We have reported that in chronic nephropatic pediatric patients there was a marked increase in plasma angiotensin-(1-7)^21^. Herein we evaluated the concentration of alamandine-(1- 5) patients in order to verify whether a similar change would be present for alamandine and alamandine-(1-5). A marked increase in alamandine-(1-5) was observed in nephrotic syndrome patients (fig. 11C).

## DISCUSSION

Using multiple approaches, we were able to demonstrate that alamandine-(1-5) is a biologically active end-product of the renin-angiotensin system. Evidence was also obtained for the presence of this pentapeptide in the human and rodent circulation, suggesting that it may take part of the complex modulatory role of the RAS in the body. In addition, a pronounced effect of alamandine-(1-5) in the heart contractility was unmasked in this study suggesting a potential role of this peptide in the heart function modulation.

Contrasting to the vasorelaxing effect of alamandine in aortic rings, a contractile effect was produced by alamandine-(1-5) in this preparation. Considering that there were no important changes in the contractile effect in endothelium denuded vessels or by pre-treatment with L-Name, a direct effect on vascular smooth muscle would be possible. Indeed presence of Mas, MrgD and AT2R, receptors probably involved in this alamandine-(1-5) effect (see below), was reported in VSM cells ^22–26^. It should be pointed out that, in view of the absence of losartan effect on the alamandine-(1-5) effect in aortic rings, AT1R appears not to play a role in the alamandine- (1-5) actions in mice aortic rings. Contrasting to aortic rings, in mesenteric arteries the pentapeptide produced vasorelaxation. Actually, a differential effect of agonists in aortic rings as opposed to other vessels is a usual finding ^27^. NO release is usually a major player in small vessels while prostanoides are reported to be also involved in mouse aortic rings preparations ^27^.

Another key finding of this study was the pronounced effect of alamandine-(1-5) in the isolated heart contractility. A finding confirmed at the cellular level by using the isolated adult ventricular myocyte preparation. The pentapeptide induced a marked decrease in cardiomyocyte contractility, and speed parameters. The reduction in the Ca^2+^ transient observed in cells exposed to alamandine-(1-5) further confirms the contractility findings and provides the mechanistic basis, at least in part, for the reduced contraction. In keeping with the dramatic effects observed in isolated ventricular myocytes, alamandine-(1-5) produced marked changes in the heart function in isolated rat heart preparation. In concentrations as low as 20 pmol.L^-1^ the peptide decreased the inotropism as evidenced by a decrease in LEVSP, dP/dt_max_ and dP/dt_min_. In addition at the lower concentration (20 pmol.L^-1^) the peptide produced a mild progressive and sustained increase in the perfusion pressure, suggesting a slight coronary vasoconstriction. At a 10-fold higher concentration the effects of alamandine-(1-5) were similar to those observed with the smaller concentration but mostly transients, suggesting dessensitization or involvement of other mechanisms. Despite the obvious need of future studies, the findings in isolated hearts reinforced the observations made in isolated cardiomyocytes, suggesting that this peptide could be a unique potent modulator of heart function.

One of the most intriguing aspects of our study is the fact that alamandine-(1-5) appears to be an agonist for all know receptors of the alternative arm of the RAS (Mas, MrgD ant AT2R) as evidenced in CHO cells expressing the different receptors. A new level of complexity is added by the findings in aortic rings. In this preparation, rings taken from MrgD or AT2R deficient mice had a marked reduction of the response to the peptide. While in rings from MAS deficient mice the response to alamandine-(1-5) was unaltered. This observation raises many possibilities including oligomerization ^28^, signaling pathways dependents on a critical common downstream component and allosteric modulation among others. One may argue that the involvement of multiple GPCR in the effects of alamandine-(1-5) indicates non-specificity, however its contractile action in aortic rings was not influenced by the AT1R antagonist Losartan alone or in rings from Mas-KO mice. In endothelial cells (EA.hy926) acute treatment with alamandine-(1-5) increased NO production. Several studies have provided evidence that Ang-(1-7) in the central nervous system enhances the baroreceptor reflex control of heart rate, especially its bradycardic component, while Ang II attenuates it ^29–31^. Besides, acute ICV infusion of Ang-(1-7) also improves baroreflex in hypertensive animals ^32–34^, while central infusion of a selective Ang-(1-7) antagonist blocks the improvement in baroreflex function produced by Ang-(1-7) in SHR ^34^. Arterial baroreceptor dysfunction is a well-known factor involved in increased blood pressure variability, which has been closely correlated to end-organ damage ^35^. Central infusion of alamandine-(1-5) in SHR improves baroreflex control, even with the maintenance of high-pressure levels. Improvement in baroreflex control has a direct impact on the autonomic control of HR, thus we also evaluated the autonomic drive to the heart through pharmacological blockade. Interestingly, central infusion of alamandine-(1-5) did not produce significant changes in cardiac autonomic balance both in normotensive or hypertensive strains. These findings suggest, at least for acute treatment, that the improvement in baroreflex could be related to a direct effect of alamandine-(1-5) in neuronal circuitry sensibility within the brain baroreceptor pathway.

The insular cortex can be broadly divided into two regions: the anterior insula and the posterior insula. Each region contributes to a distinct set of processes. The rostral IC is heavily involved in interoception, the processing of internal bodily signals like hunger, thirst, pain and cardiovascular control as previously described by our group. It also plays a crucial role in emotion processing, particularly in integrating emotional experiences with bodily sensations, decision-making and self-awareness. Recent discoveries highlight a surprising level of specialization within the insula, suggesting it is responsible for specific peptides, such as alamandine ^3^. The fact that the alamandine product, alamandine-(1-5), displayed a selective effect, increasing blood pressure without influencing HR or renal nerve activity is intriguing and deserves further studies. While the exact mechanisms are not understood, this newfound ínsula selective stimulator opens exciting avenues for further research on the role of alatensins ^36^ in this region.

We have observed a potent and long-lasting (>6 h) anti-hypertensive effect following administration of a single dose (30 ug/Kg) of alamandine-(1-5) in SHR. A similar long-lasting anti-hypertensive effect was previously observed for alamandine, angiotensin-(1-7) and BPPs peptides in our laboratory ^2,37,38^. This suggests that these peptides upon binding to their receptors/targets could trigger a long-lasting signaling mechanism which influences vascular tonus and heart function. Whether this is selective for SHR or could be present in other hypertension models needs to be clarified.

The major finding in the echocardiography in SHR was a 33% decrease in cardiac output due to a significant decrease in stroke volume. This change probably contributed to the anti- hypertensive effect of alamandine-(⅖) in these animals. On the other hand, a large variation in ejection fraction and fractional shortening was observed, precluding a contribution of a direct effect of the peptide in these variables. It is very difficult to compare the changes in mice isolated cardiomyocyte and isolated perfused hearts with the effects of the peptide in vivo in SHR, however it is clear from our results that a direct or indirect alteration in heart function is also present in SHR following administration of alamandine-(1-5).

To investigate the formation of this pentapeptide, we assessed its generation from alamandine using plasma samples from Angiotensinogen-KO mice to eliminate potential confounding factors. There was a rapid formation of alamandine-(1-5) in the plasma incubated with alamandine. However, in plasma incubated with alamandine in the presence of captopril the formation of alamandine-(1-5) was suppressed, suggesting a significant involvement of ACE in its formation from Alamandine. Our study did not explore the potential formation of alamandine- (1-5) from Angiotensin-(1-5) through decarboxylation of the aspartate residue to alanine. This intriguing possibility warrants investigation in future studies. Of note, we detected circulating alamandine-(1-5) in plasma samples obtained from healthy human volunteers (Figure 11A) and C57Bl6/J mice (Figure 11B). More important, our present data about the concentration of alamandine-(1-5) plasma levels showed a marked elevation in pediatric patients with nephrotic syndrome in comparison to healthy controls. The precise meaning of this finding is currently unknown and our sample size is limited. However, we might speculate that, as previously reported by other RAS components^39^, alamandine-(1-5) may have a role in the pathophysiology of primary nephrotic syndrome. These novel findings combined with the many effects of alamandine-(1-5) unmasked in our study, indicates that this pentapeptide may be involved in several actions of the RAS.

It is now considered that the RAS comprises two main axes, classical and alternative, which in most circumstances have opposite actions in many organs and systems. Our current findings suggest that the RAS complexity might be far beyond this current view. In this regard, alamandine-(1-5) which is a novel end-product of the vasodilator axis has indeed many selective actions that differentiate it from our current understanding of this axis. Furthermore, our data suggest that this pentapeptide circulates in the human blood and might be importantly involved in the modulation of heart inotropism by possibly influencing Ca^+2^ homeostasis. Our current data also suggests that the actions of alamandine-(1-5) are mediated through a complex interaction of Mas, MrgD and AT2 receptors.

In conclusion, in this study we identified and characterized a new component of the RAS, alamandine-(1-5), which may be importantly involved in cardiac function modulation and kidney pathophysiology.

