## supplementary data for "Identification and characterization of alamandine-(1-5), a new component of the Renin-Angiotensin System with unique properties"

Supplementary Methods

Evaluation of cardiac autonomic control

The cardiac autonomic tone was determined by pharmacological blockade of either the sympathetic or parasympathetic tone to the heart in two sequential days in a reversed order. On the 1st day, 30 min after the baroreflex test, rats received methylatropine, a muscarinic blocker (3 mg/kg i.v.) and the maximum HR was recorded. Between 15 to 20 minutes later, animals received propranolol, a β-adrenergic blocker (4 mg/kg i.v.) and intrinsic HR (iHR) was recorded. Twenty-four hours later, these blockers were injected in the reverse order to obtain the minimal HR, after propranolol injection, and the iHR of the 2nd day. The iHR was considered to be the heart rate after complete autonomic blockade i.e. muscarinic cholinergic and a β-adrenergic receptor blockade, and calculated as the average of the values recorded at the 1st and 2nd day. The sympathetic tonus was calculated by the difference between the maximum HR (after methylatropine on the 1st day) and intrinsic HR, and the parasympathetic tonus, by the difference between intrinsic HR and the minimum HR (after propranolol on the 2nd day) (1).

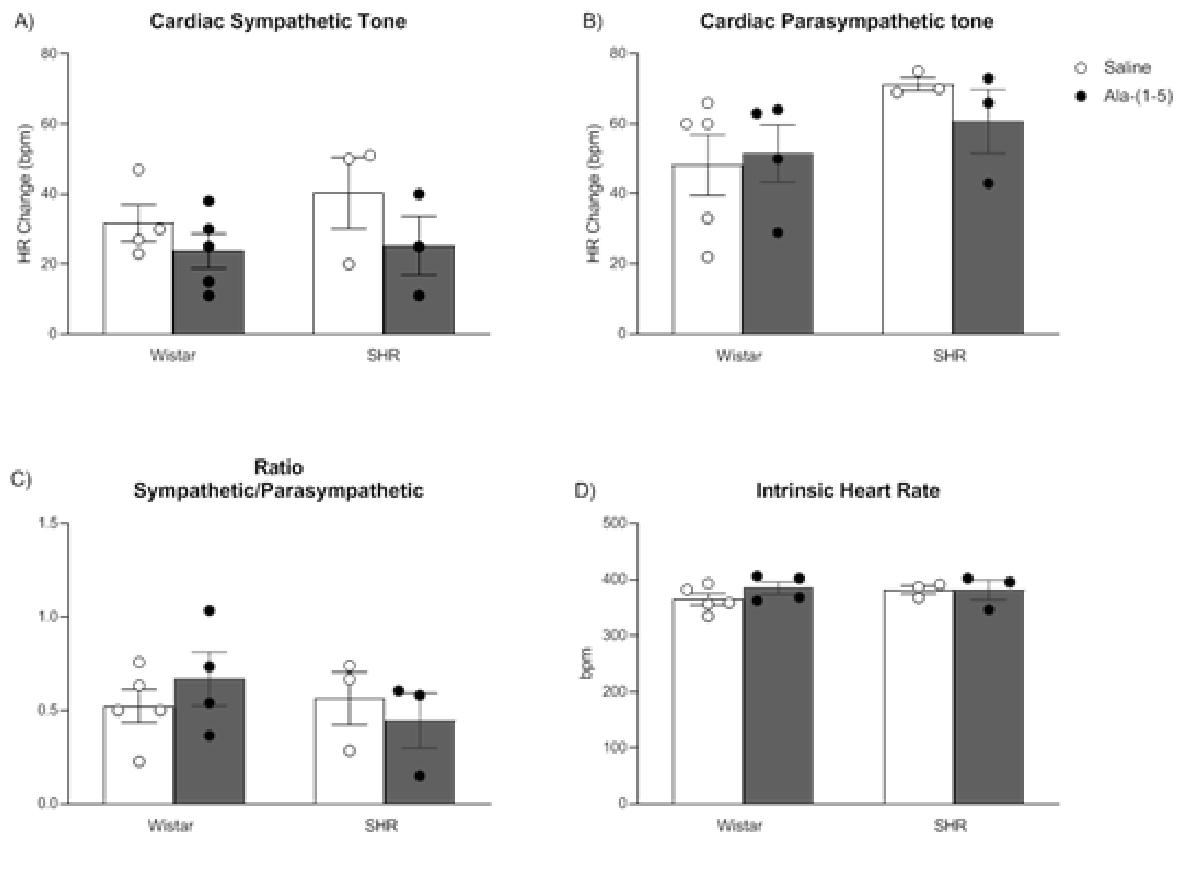

**Figure S1**. Cardiac autonomic tone changes to alamandine-(1-5). Cardiac sympathetic tone (HR Change, bpm; A), cardiac parasympathetic tone (HR Change, bpm; B), ratio of cardiac sympathetic and parasympathetic tone (C) and intrinsic heart rate (bpm; D) after 90 minutes of ICV infusion (12 µl/hour) of saline or alamandine-(1-5) [(Ala-(1-5); 0.04 µg/µl] in Wistar and SHR rats. Values are mean ± S.E.M.

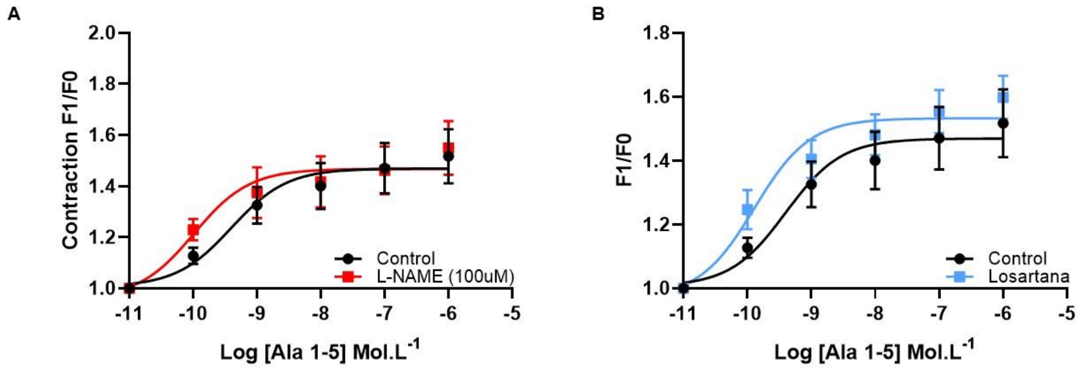


**Figure S2.** Effects of L-NAME and losartan on the alamandine-(1-5) in isolated aortic rings. A. Cumulative concentration of alamandine-(1-5) (Ala-(1-5)) in aortic rings pre-treated with L-NAME 100 µmol. L-1; B. Cumulative concentration of alamandine-(1-5) (Ala-(1-5)) in aortic rings pre-treated with losartan10-6 mol. L-1. Data were expressed as mean ± SEM and analyzed by two-way ANOVA followed by Fisher's multiple comparisons test. P<0.05.

Table 1: Demographic characteristics of patients with primary nephrotic syndrome and healthy controls matched by age and gender.

| **Demographics** | **Patients (n = 6)** | **Controls (n = 6)** | **P value** |
|---|---|---|---|
| **Age (months)** | 81.00±49.31 | 82.12±33.24 | 0.86 |
| **Sex (male)** | 3 (50%) | 3 (50%) | - |
| **Ethnicity** | | | |
| **White** | 4 (66.7%) | 4 (66.7%) | - |
| **Black** | 2 (33.3%) | 2 (33.3%) | - |
| **Weight (kg)** | 25.53±17.20 | 26.74±19.58 | 0.78 |
